# The chromatin organization of a chlorarachniophyte nucleomorph genome

**DOI:** 10.1101/2021.09.01.458626

**Authors:** Georgi K. Marinov, Xinyi Chen, Tong Wu, Chuan He, Arthur R. Grossman, Anshul Kundaje, William J. Greenleaf

## Abstract

Nucleomoprhs are remnants of secondary endosymbiotic events between two eukaryote cells wherein the endosymbiont has retained its eukaryotic nucleus. Nucleomorphs have evolved at least twice independently, in chlorarachniophytes and cryptophytes, yet they have converged on a remarkably similar genomic architecture, characterized by the most extreme compression and miniaturization among all known eukaryotic genomes. Previous computational studies have suggested that nucleomorph chromatin likely exhibits a number of divergent features. In this work, we provide the first maps of open chromatin, active transcription, and three-dimensional organization for the nucleomorph genome of the chlorarachniophyte *Bigelowiella natans*. We find that the *B. natans* nucleomorph genome exists in a highly accessible state, akin to that of ribosomal DNA in some other eukaryotes, and that it is highly transcribed over its entire length, with few signs of polymerase pausing at transcription start sites (TSSs). At the same time, most nucleomorph TSSs show very strong nucleosome positioning. Chromosome conformation (Hi-C) maps reveal that nucleomorph chromosomes interact with one other at their telomeric regions, and show the relative contact frequencies between the multiple genomic compartments of distinct origin that *B. natans* cells contain.

## Introduction

Endosymbiosis, especially between a eukaryotic host and a prokaryote, is a common event in the evolution of eukaryotes, and subsequent changes in the host and endosymbiont genomes often follow similar general trends. One such trend is the reduction of the endosymbiont’s genome due to gene loss and endosymbiotic gene transfer^1,2^ (EGT) into the host’s nucleus, the classic example of which are the extremely reduced genomes of plastids and mitochondria that evolved as the bacterial progenitors of these organelles underwent organellogenesis. This trend is also strongly manifested in the fate of secondary endosymbionts (eukaryotes that become endosymbionts of other eukaryotes). Such endosymbiotic events have occurred on multiple occasions in the evolution of eukaryotes^3^, usually resulting in retention of the plastid of the photosynthetic eukaryotic endosymbiont (as a secondary plastid) while the nucleus of the endosymbiont is lost entirely. However, several notable exceptions to this general rule do exist. One is the dinotoms, the result of an endosymbiosis between a dinoflagellate host and a diatom, in which the diatom has not been substantially reduced^4,5^. More striking are the nucleomorphs, which are best known from the chlorarachniophytes and the cryptophytes (but may in fact have arisen in other groups too, such as some dinoflagellates^6,7^). Nucleomorphs retain a vestigial nucleus with a highly reduced but still functional remnant of the endosymbiont’s genome^8,9^.

A remarkable feature of chlorarachniophyte and cryptophyte nucleomorphs is that they have evolved independently, from a green and a red alga, respectively, yet their genomes exhibit surprisingly convergent properties^10,11^. In both cases, the genomes of their nucleomorphs are the smallest known among all eukaryotes, usually just a few hundred kilobases in size (~380 kbp for the chlorarachniophyte *B. natans*). All sequenced nucleomorph genomes are organized into three highly AT-rich chromosomes, in which arrays of ribosomal RNA genes form the subtelomeric regions. These genomes are also extremely compressed, exhibiting very little intergenic space between genes, with genes even overlapping on occasions. The genes themselves are also often shortened^12–18^.

A number of important questions about the biology associated with the extremely reduced nucleomorph genome remain unanswered, including the extent of conservation and divergence of chromatin organization and transcriptional mechanisms of these extremely reduced nuclei relative to that of a convention eukaryotic genome. Previous computational analysis of nucleomorph genome sequences^19^ has suggested that a considerable degree of deviation from the conventional eukaryotic state is likely to have developed in nucleomorphs. For example, histone proteins are ancestral to all eukaryotes, and the key posttranscriptional modifications (PTMs) that they carry also date back to the last eukaryotic common ancestor (LECA) and are extremely conserved in nearly all branches of the eukaryotic tree^20^, with the notable exception of dinoflagellates^21^. This is likely because these PTMs are deposited in a highly regulated manner on specific residues of histones, and are then read out by various effector proteins, thus playing crucial roles in practically all aspects of chromatin biology, such as the regulation of gene expression, the transcriptional cycle, the formation of repressive heterochromatin, mitotic condensation of chromosomes, DNA repair, and many others (in what is often referred to as “histone code”^22^).

Nucleomorphs appear to be one of the few^19,21^ exceptions to this general rule. Inside nucleomorph genomes, in both chlorarachniophytes and cryptophytes, only two histone genes are encoded, one for H3 and for H4, with H2A and H2B encoded by the host nuclear genome and imported from the host’s cytoplasm^23^. Sequence analyses of the H3 and H4 proteins show remarkable divergence from the typical amino acid sequence in eukaryotes; specifically, the chlorarachniophyte histones have lost nearly all key histone code residues^19^. Furthermore, the heptad repeats in the C-terminal domain (CTD) tail of the Rpb1 subunit of RNA Polymerase II, which are highly conserved in eukaryotes^24^ and key to the their transcriptional cycle and mRNA processing^25^, have also been lost.

These observations suggest that the nucleomorph chromatin and chromatin-based regulatory mechanisms may be unconventional compared to those of other eukaryotes. For example, nucleomorphs may organize and protect DNA differently than other eukaryotes, nucleomorph promoters may display atypical signatures of nucleosome depletion and positioning, histone modifications, etc., and relation of these marks to transcriptional activity, or they may exhibit unique 3D genomic organization. However, none of these features associated with nucleomorph chromatin or gene expression regulation has been directly studied.

In this work we map chromatin accessibility, active transcription, and three-dimensional (3D) genome organization in the chlorarachniophyte *Bigelowiella natans* to address these gaps in our knowledge of nucleomorph biology. We find that nucleomorph chromosomes exist in a highly accessible state, reminiscent of what is observed for ribosomal DNA (rDNA) in other eukaryotes, such as budding yeast, where rDNA is thought to be fully nucleosome-free when actively transcribed^26–28^. However, nucleomorph promoters are associated with strongly positioned nucleosomes, and they exhibit a distinct nucleosome-free region upstream of the transcription start site (TSS). Active transcription is nearly uniformly distributed across nucleomorph genomes, with the exception of elevated transcription and chromatin accessibility at the subtelomeric rDNA genes. We find few signs of RNA polymerase pausing over promoters. Nucleomorph chromosomes form a network of telomere-to-telomere interactions in 3D space, and also fold on themselves, but centromeres do not preferentially interact with each other. Curiously, the genome of the *B. natans* mitochondrion, which derives from the host, exhibits an elevated Hi-C *trans* contact frequency with the genomes of the endosymbiont compartments (the plastid and the nucleomorph) than it does with the host genome. These results provide novel insights into chlorarachniophyte nucleomorph chromatin structure and a framework for future mechanistic studies of transcriptional and regulatory biology in nucleomorphs.

## Results

### Chromatin accessibility in nucleomorphs

To study the chromatin structure of the *B. natans* nucleomorph genome, we carried out ATAC-seq experiments in *B. natans* grown under standard conditions (see Methods). As *B. natans* has four different genomic compartments (Figure 1A) – nucleus, nucleomorph, mitochondrion and plastid – we first examined the fragment length distribution in each (Figure 1B). The nucleus exhibits a subnucleosomal peak at ~ 100 bp as well as a second, most likely nucleosomal, peak (or a “shoulder” in the curve) at ~200 bp. In contrast, the nucleomorph displays two peaks, one at ≤100 bp and another at ~220 bp, which are tentatively interpreted as subnucleosomal and a nucleosomal one (see further below for a more detailed discussion). The mitochondrion and the plastid fragment length distributions are unimodal, consistent with the open DNA structure expected from these compartments which do not contain nucleosomes.

**Figure 1:**
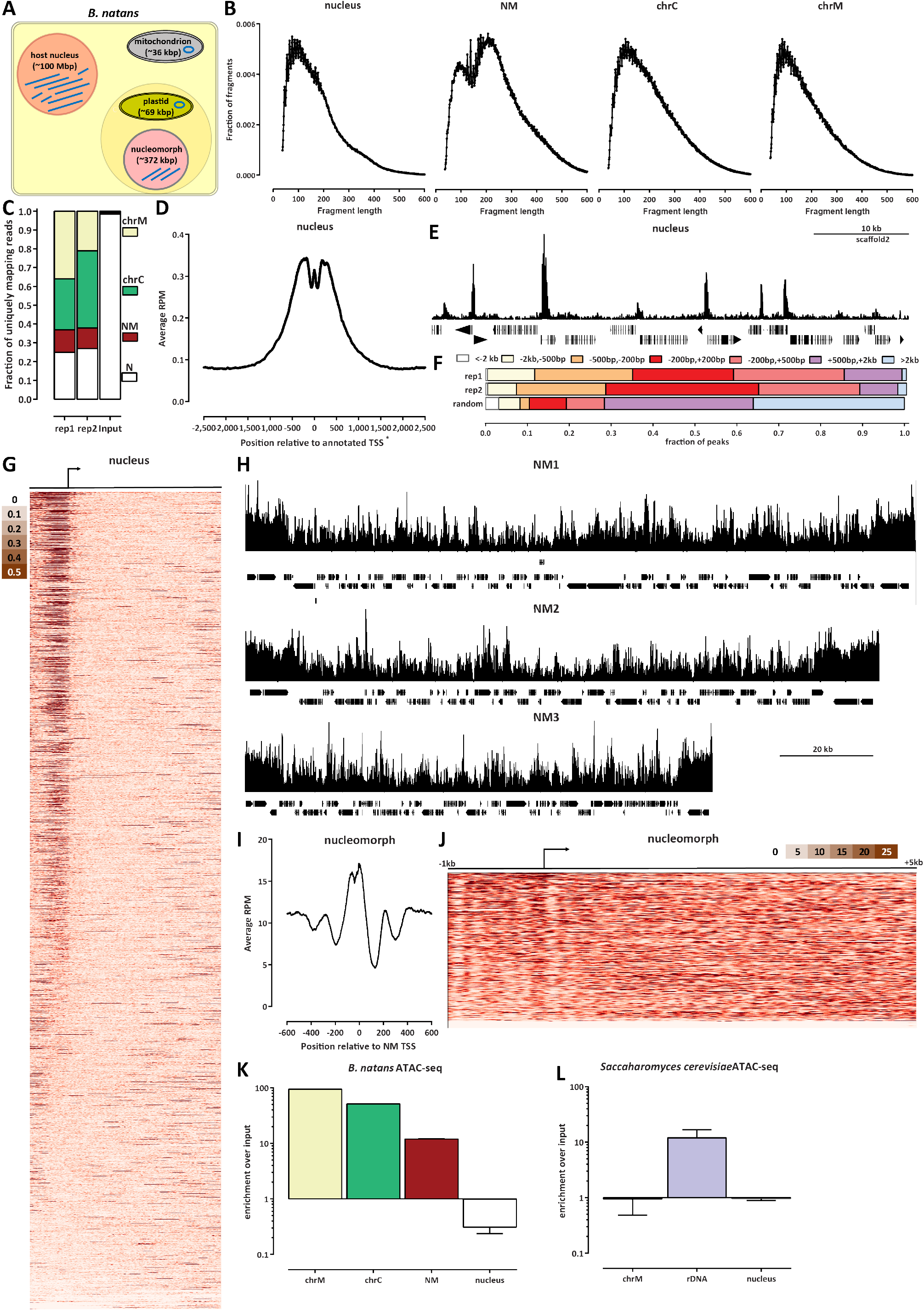
The chromatin accessibility landscape of the *B. natans* nuclear and nucleo-morph genomes. (A) Schematic outline of the different genomic compartments in a *B. natans* cell. (B) ATAC-seq fragment length distribution in the different genomic compartments. (C) Distribution of mapped ATAC-seq reads across genomic compartments. (D) ATAC-seq read coverage metaplot around nuclear TSSs. (E) Snapshot of an ATAC-seq profile at a typical nuclear locus. (F) Distribution of ATAC-seq called peaks in the nucleus relative to TSSs. The “random” distribution was generated by splitting the genome in 500-bp bins and taking the boundary coordinates of each bin as “peaks”. (G) ATAC-seq profiles around all nuclear genes. (H) ATAC-seq profiles over the NM1, NM2 and NM3 nucleomorph chromosomes. (I) ATAC-seq read coverage metaplot around nucleomorph TSSs. (J) ATAC-seq profiles around all nucleomorph genes. (K) The nucleomorph genome is ~10× enriched in ATAC-seq datasets relative to the nuclear genome. Shown is the ratio of normalized mapped ATAC-seq peaks for each of the compartments relative to the normalized mapped reads in an input sample (a Hi-C dataset mapped in a single-end format). (L) Nucleomorph accessibility is comparable to the accessibility of rDNA loci in the budding yeast *S. cerevisiae*, which exist in a fully nucleosome-free conformation when expressed.

We then examined the distribution of reads across the compartments (Figure 1C). As expected from the lack of nucleosomal protection over mitochondrial and plastid DNA, *B. natans* ATAC libraries are dominated by reads mapping to those compartments. However, curiously, nucleomorph-mapping reads also represented a much larger fraction of mapped reads than expected from the portion of genomic real estate that the nucleomorph genome comprises, and also relative to what is seen in input samples, suggesting that the nucleomorph might exist in a preferentially accessible chromatin state.

We next turned our attention to ATAC-seq profiles in the nucleus, both to characterize accessibility in the *B. natans* host genome, and to verify the quality of the ATAC-seq libraries. Figure 1D shows the average ATAC-seq signal over annotated *B. natans* TSSs; it is enriched over promoters, as expected from successful ATAC-seq experiments (we note that the shape of the metaplot is somewhat distorted by the fact that available annotations do not actually include the actual TSSs, but only the sites of translation initiation, with most 5’UTR missing). Examination of browser tracks confirmed the enrichment over TSSs (Figure 1E), and did not reveal obvious open chromatin sites outside promoters. We carried out peak calling using MACS2^29^, and the distribution of called peaks was also strongly centered on promoters, with almost no open chromatin regions outside the ±2 kbp range around TSSs. Thus *B. natans* appears to have a functional genomic organization typical for a eukaryote with a small compact genome such as yeast, with all regulatory elements located immediately adjacent to TSSs, and few to no distal regulatory elements that exhibit increased accessibility. In addition, in standard *B. natans* culture conditions, the majority of promoters exhibit an open chromatin configuration (Figure 1G).

Genome browser examination of ATAC-seq profiles over the nucleomorph genome (Figure 1H) showed high levels of chromatin accessibility throughout all chromosomes, with numerous localized peaks and generally increased accessibility over the rDNA located near telomeres. Strikingly, the average ATAC-seq profile over nucleomorph TSSs (Figure 1I) showed a strong increase in accessibility around the TSS, but also a clear signature of multiple positioned nucleosomes around each TSS (a clear +1 nucleosome immediately downstream of the TSS, as well as a putative +2 one, together with a −1 nucleosome upstream of the TSS). This phasing is also clearly visible from the individual ATAC-seq profiles over each nucleomorph gene (Figure 1J).

We then quantified the extent of increased accessibility over organellar genomes by calculating the enrichment of ATAC-seq signal relative to the total DNA mass as measured by an input sample. We find that the nucleomorph is ~10× enriched in ATAC-seq libraries, compared to ~100× and ~50× for the mitochondrion and plastid genomes, respectively (Figure 1K). Notably, this enrichment is comparable to what is observed for rDNA genes in the budding yeast *S. cerevisiae* (Figure 1L), which are known to exist in an almost fully nucleosome-free configuration when actively transcribed, which is thought to be ~50% of the time^26–28,30^.

Thus the nucleomorph apparently exists in a highly accessible state. Of note, this estimation is not driven by the rDNA genes within it, although those are indeed more accessible than the rest of the nucleomorph genome, as the difference in accessibility between the rDNA arrays and the rest of the genome is on the order of ~2× and they occupy a minor (~11%) portion of it.

However, nucleomorph TSSs show very strong nucleosome positioning. To more accurately analyze nucleosome positioning in both the nuclear and the nucleomorph compartments, we applied the NucleoATAC algorithm^31^ over the whole nucleomorph genome and over the 1-kb regions centered on annotated 5’ gene ends in the nucleus. We identified 7,251 and 1,440 positioned nucleosomes in the nucleus and in the nucleomorph, respectively. The distribution of the nuclear nucleosomes peaked shortly downstream of TSSs (Figure 2A), suggesting that nuclear TSSs are also associated with a positioned +1 nucleosome. A V-plot^32^ analysis showed that the ATAC-seq fragment lengths associated with these nucleosomes are in the 175-200 bp range, and that subnucleosomal fragments are located in the immediate vicinity (Figure 2A). In contrast, in the nucleomorph we observe three nucleosomes positioned in the vicinity of the TSS (+1, +2, and −1; Figure 2C), but ATAC-seq fragment lengths associated with these nucleosomes are larger, in the 200-225 bp range (Figure 2D).

**Figure 2:**
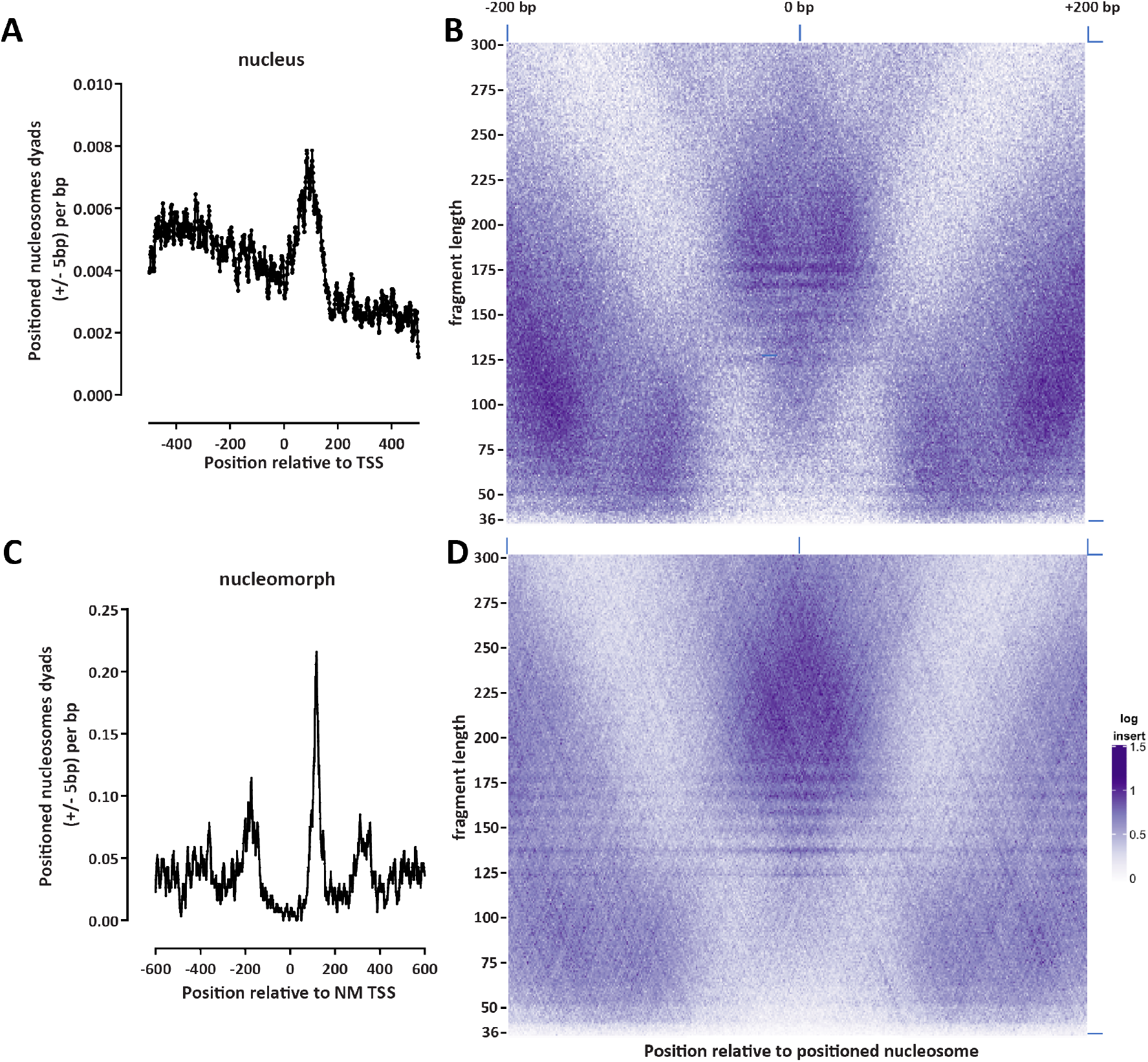
Nucleosome positioning in the *B. natans* nuclear and nucleomorph genomes. (A) Location of positioned nucleosomes (determined by NucleoATAC) relative to annotated TSSs in the *B. natans* nucleus (shown are dyad positions extended by ±5 bp). (B) V-plot of ATAC-seq fragment distribution around positioned nucleosomes in the nucleus. (C) Location of positioned nucleosomes (determined by NucleoATAC) relative to annotated TSSs in the *B. natans* nucleomorph (shown are dyad positions extended by ±5 bp). (D) V-plot of ATAC-seq fragment distribution around positioned nucleosomes in the nucleomorph.

### Transcriptional activity in the nucleomorph genome

Next, we studied the patterns of active transcription in the nucleomorph. To this end, we deployed the KAS-seq assay^33^, which maps single-stranded DNA (ssDNA) by specifically labeling unpaired guanines with N_3_-kethoxal, to which biotin can then be attached using click chemistry, allowing for regions containing ssDNA to be specifically enriched. Most ssDNA in the cell is usually associated with RNA polymerase bubbles^33^, thus KAS-seq is a good proxy for active transcription.

In the *B. natans* nucleus, KAS-seq shows enrichment over promoters and over actively transcribed genes (Figure 3A-B), as expected based on patterns observed in other eukaryotes^33^, indicative of RNA polymerase spending more time near the TSS. However, we observe only very weak correlation between promoter accessibility and active transcription (Supplementary Figure 1), suggesting significant decoupling between the opening of nucleosome depleted promoter-proximal regions and the regulation of active transcription in *B. natans*.

**Figure 3:**
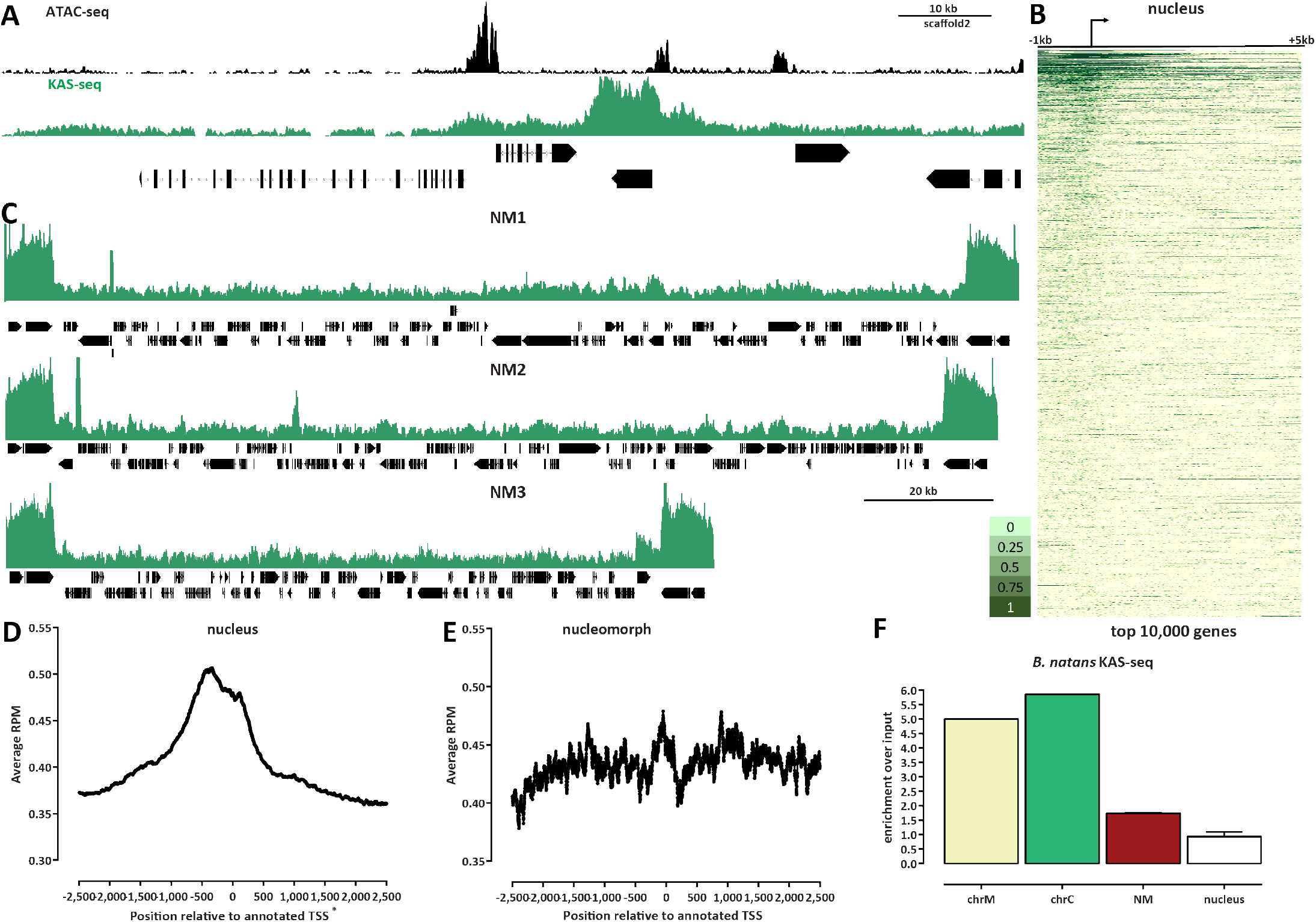
The active transcription landscape of the B. *natans* nuclear and nucleomorph genomes as measured by KAS-seq. (A) KAS-seq and ATAC-seq profiles at a typical nuclear locus. (B) KAS-seq profiles over the top 10,000 (by KAS signal) nuclear genes. (C) KAS-seq profiles over the NM1, NM2 and NM3 nucleomorph chromosomes. (D) Average KAS-seq profile over nuclear gene TSSs. (E) Average KAS-seq profile over nucleomorph TSSs.s (F) Relative enrichment of KAS-seq signal in the different *B. natans* genomic compartments. Shown is the ratio of normalized mapped KAS-seq peaks for each of the compartments relative to the normalized mapped reads in an input sample (a Hi-C dataset mapped in a single-end format).

In the nucleomorph, we see largely uniform levels of KAS-seq signal, with the exception of the rDNA genes, and three localized peaks, one on the first, and two on the second nucleomorph chromosomes (Figure 3C-E). The increased transcription of the rDNA genes is consistent with their higher accessibility observed in ATAC-seq data. We quantified the overall enrichment of active transcription in the different compartments and found that the nucleomorph is ~2-fold enriched in KAS-seq datasets than the nucleus (Figure 3F) relative to an input sample.

These observations, based on measuring actual active transcription, corroborate previous reports, based on transcriptomic analysis, of high, pervasive, and largely uniform transcriptional activity over most of the nucleomorph genome^34–37^. However, rDNA genes were removed in some of these analyses^34^ while we identify them as a transcriptional unit existing in a distinct state from the rest of the nucleomorph genome (in the analysis presented here, multimapping reads were retained and normalized, allowing us to measure accessibility and transcription levels over the rDNA genes; see the Methods section for more details).

### Three-dimensional organization of the *B. natans* nucleomorph genome

Finally, we mapped the three-dimensional genome organization in *B. natans* using *in situ* chromosomal conformation capture (Hi-C^38^). We employed a modified protocol for the highly AT-rich nucleomorph genomes (see Methods for details) and generated high-resolution 1-kbp maps, which allow us to investigate the fine features of the small nucleomorph chromosomes.

Hi-C maps reveal that the nucleomorph chromosomes often exist in a folded conformation, in which the two chromosome ends contact each other (Figure 4A-B). In addition, the telomeric regions of all nucleomorph chromosomes physically associate with each other, forming a telomeric network of interactions (Figure 4A). In many eukaryotes, a centromeric interaction network is also observed^39^, but enriched interchromosomal interactions in nucleomorphs appear to be only telomeric. We do not observe much internal structure inside individual nucleomorph chromosomes, with the exception of NM2, in which one potential loop interaction is seen; its mechanistic origins are currently unclear as its singular nature prevents the identification of sequence drivers of its formation.

**Figure 4:**
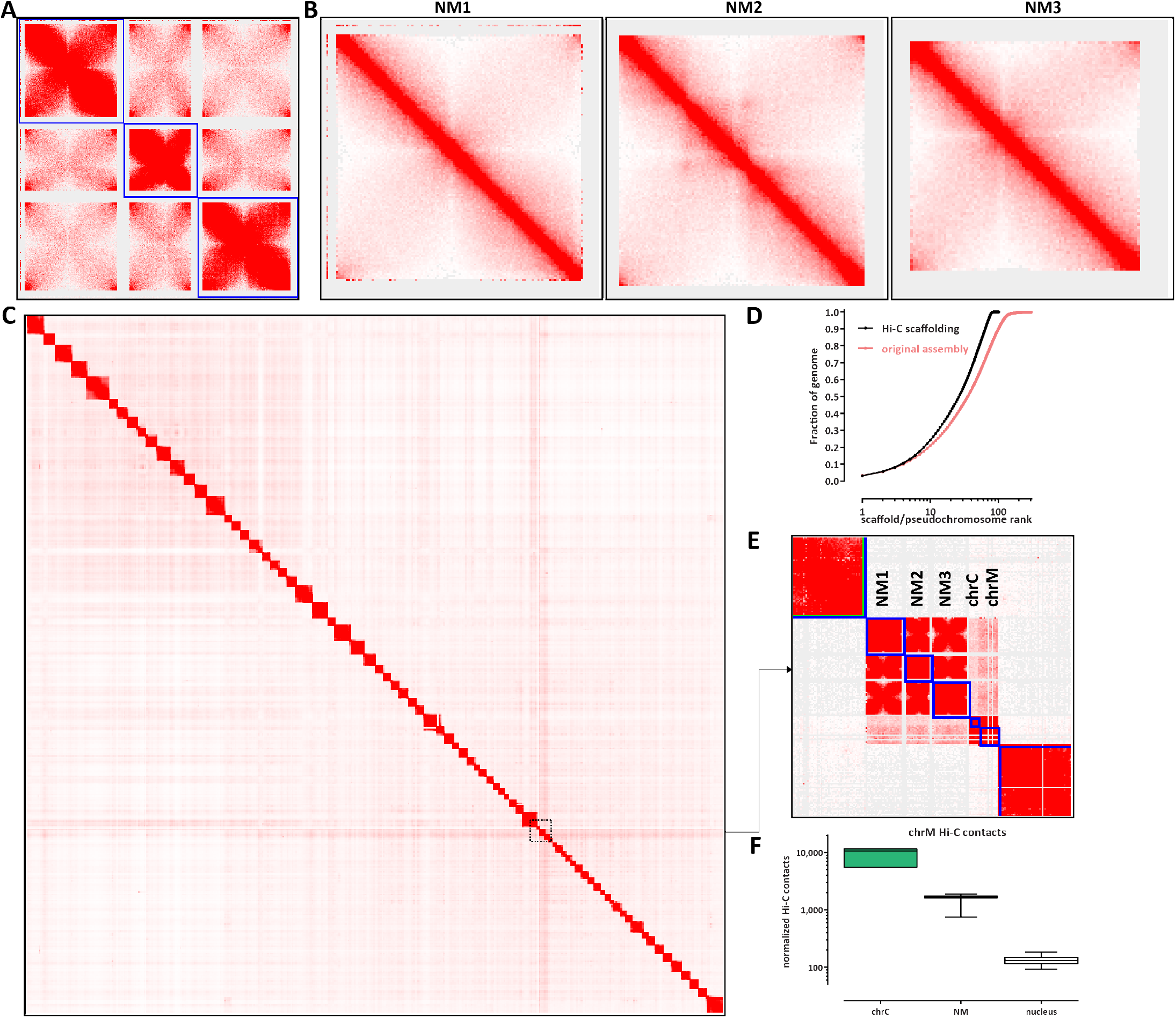
Three-dimensional organization of *B. natans* nucleomorph chromosomes. (A) Hi-C maps (5-kbp resolution) of the three NM chromosomes reveals a network of telomere-to-telomere interactions as the main 3D organizational feature of the nucleomorph. (B) High-resolution (1-kbp) maps of the individual NM chromosomes. (C) and (D) Global scaffolding of the B. *natans* genome. (E) and (F) The B. *natans* mitochondrion exhibits higher Hi-C *trans* contacts with the endosymbiont compartments than with the nucleus.

We also used our Hi-C data to generate a chromosome-level scaffolding^40^ of the existing assembly of the *B. natans* nuclear genome^41^, which originally consisted of 302 nuclear contigs. Our chromosome-level assembly identifies 79 pseudochromosomes; the smallest is ~ 350 kbp, and the largest is ~3 Mbp. This assembly retains only 18 smaller unplaced contigs, the largest being 8,753 bp (Figure 4D).

We made one surprising observation when manually finalizing the chromosome-level assembly – although the mitochondrion is topologically derived from the host (Figure 1A) and is separated from the nucleomoprh and plastid genomes in the endosymbiont by several membranes, it exhibits elevated Hi-C *trans* contacts with both plastid and nucleomorph chromosomes. This preferential enrichment can be visually seen in the Hi-C maps themselves (Figure 4E), and was also confirmed by a systematic analysis of chrM *trans* contacts with all other chromosomes (Figure 4F). We also note that we obtain the same result with all available methods for normalizing Hi-C data (Supplementary Figure 2). While both the plastid and the mitochondrial genomes exist in high copy numbers, the nucleomorph genome has the same copy number as the nuclear genome (as shown by our input samples, in which read coverage over the nucleomorph is the same as read coverage over the nucleus), thus it is likely that these preferential Hi-C contacts likely indeed represent frequent physical proximity in the cell, which then leads to ligation events with nucleomorph chromosomes during the *in situ* Hi-C procedure.

## Discussion

This study presents the first analysis of physical chromatin organization in a nucleomorph genome, in the chlorarachniophyte *B. natans*, using a combination of ATAC-seq, Hi-C, and KAS-seq measurements. We also provide a near-complete chromosome-level scaffolding of the nuclear genome by taking advantage of the physical proximity information provided by Hi-C data, and assess the extent of physical interactions between the different genomic compartments.

While it was previously suspected that nucleomorphs are very highly transcriptionally active, we demonstrate that this activity is also reflected at the level of chromatin structure, as nucleomorph chromosomes are much more highly accessible than those in the nucleus. Previous transcriptomic analyses also suggested pervasive largely uniform transcription levels that also do not change much between conditions^34,35,37^, and this is also what is seen at the level of the measurements of active transcription by KAS-seq, with the notable exception of the rDNA genes, which are much more strongly transcribed than the rest of the nucleomorph (and also exhibit elevated accessibility). Taken together, these results suggest the possibility of limited transcriptional regulation in the nucleomorph (which may also be related to the strong divergence of nucleomorph histones H3 and H4 and the absence in them of most key residues involved in regulatory functions). However, nucleomorph promoters exhibit a very prominent upstream nucleosome depleted region and strong degree of nucleosome positioning. How this promoter architecture is generated by sequence elements associated with each promoter is at present not known. It also remains opaque whether these elements merely indicate the location of transcription initiation or if sequence elements with regulatory activity can influence the levels of transcription. To dissect the function of these elements, methods for the direct genetic manipulation of nucleomorphs will be needed. Somewhat surprisingly, this strong nucleosome positioning at TSSs is not associated with promoter pausing by the polymerase; elucidating the mechanistic details of transcription initiation and initial nucleosome clearance will likely resolve this apparent contradiction.

The presence of strongly positioned promoter-proximal nucleosomes also suggests that nucleosomes in different locations in the nucleomorph may in fact exist in distinct chromatin states, but what these might be given the lack of the classical histone posttranslational modifications in the nucleomorph histones is a mystery. There exist only limited studies of the nucleomorph proteome^42^, and the posttranslational modifications of nucleomorph histones are yet to be studied. The difference in nucleosome protection fragment lengths between the nuclear and the nucleomorph compartment suggests that the nucleomorph may also contain a distinct linker histone(s); these issues remain to be clarified in the future.

The mechanistic origins of the preferential association between mitochondria and endosymbiont compartments in Hi-C maps are not currently clear. The mitochondrial genome is enclosed by two membranes, while the endosymbiont is enveloped by two membranes, and the plastid inside it by another two^11^. Thus it is six membranes that separate mitochondrial genome from the plastid genome, and four membranes plus a nuclear membrane exist between it and the nucleomorph chromosomes. More frequent physical proximity between mitochondria and the endosymbiont in the cell is the most likely candidate explanation, as permeabilization of membranes during fixation could allow for crosslinking between chromatin in different compartments. High-resolution imaging approaches^43,44^ should be able to test this hypothesis.

Finally, it will be instructive to compare chromatin organization across the different nucleomorph-bearing groups. Nucleomorph histones in cryptophytes are considerably closer to the conventional state of most eukaryotes, and thus determining if these organisms also exhibit elevated accessibility, strong nucleosome positioning, and lack promoter polymerase pausing will be illuminating.

## Methods

### *B. natans* cell culture

*Bigelowiella natans* strain CCMP2755 starting cultures were obtained from NCMA (National Center for Marine Algae and Microbiota) and cultured in L1-Si media on a 12-h-light:12-h-dark cycle.

### ATAC-seq experiments

ATAC-seq experiments were performed following the omni-ATAC protocol^45^.

Briefly, ~1M *B. natans* cells were centrifuged at 1,000*g*, then resuspended in 500 *μ*L 1× PBS and centrifuged again. Cells were then resuspended in 50 *μ*L ATAC-RSB-Lysis buffer (10 mM Tris-HCl pH 7.4, 10 mM NaCl, 3 mM MgCl_2_, 0.1% IGEPAL CA-630, 0.1% Tween-20, 0.01% Digitonin) and incubated on ice for 3 minutes. Subsequently 1 mL ATAC-RSB-Wash buffer (10 mM Tris-HCl pH 7.4, 10 mM NaCl, 3 mM MgCl_2_, 0.1% Tween-20, 0.01% Digitonin) were added, the tubes were inverted several times, and nuclei were centrifuged at 500 *g* for 5 min at 4 °C.

Transposition was carried out by resuspending nuclei in a mix of 25 *μ*L 2× TD buffer (20 mM Tris-HCl pH 7.6, 10 mM MgCl_2_, 20% Dimethyl Formamide), 2.5 *μ*L transposase (custom produced) and 22.5 *μ*L nuclease-free H_2_O, and incubating at 37° C for 30 min in a Thermomixer at 1000 RPM.

Transposed DNA was isolated using the MinElute PCR Purification Kit (Qiagen Cat# 28004/28006), and PCR amplified as previously before^45^. Libraries were purified using the MinElute kit, then sequenced on a Illumina NextSeq 550 instrument as 2×36mers or as 2×75mers.

### KAS-seq experiments

KAS-seq experiments were performed as previously described^33^ with some modifications.

*B. natans* cells were pelleted by centrifugation at 1000 *g* for 5 minutes at room temperature, then resuspended in 500 *μ*L of media supplemented with 5 mM N_3_ -kethoxal (final concentration). Cells were incubated at room temperature for 10 minutes, then centrifuged at 1000 *g* for 5 minutes at room temperature to remove the media with the kethoxal, and resuspended in 100 *μ*L cold 1× PBS. Genomic DNA was then extracted using the Monarch gDNA Purification Kit (NEB T3010S) following the standard protocol but with elution using 85 *μ*L 25 mM K_3_BO_3_ at pH 7.0.

The click reaction was carried out by combining 87.5 *μ*L purified and sheared DNA, 2.5 *μ*L 20 mM DBCO-PEG4-biotin (DMSO solution, Sigma 760749), and 10 *μ*L 10× PBS in a final volume of 100 *μ*L. The reaction was incubated at 37°C for 90 minutes.

DNA was purified using AMPure XP beads (50 *μ*L for a 100 *μ*L reaction or 100 *μ*L for a 200 *μ*L reaction), beads were washed on a magnetic stand twice with 80% EtOH, and eluted in 130 *μ*L 25mM K_3_BO_3_.

Purified DNA was then sheared on a Covaris E220 instrument down to ~150-400 bp size.

For streptavidin pulldown of biotin-labeled DNA, 10 *μ*L of 10 mg/mL Dynabeads MyOne Streptavidin T1 beads (Life Technologies, 65602) were separated on a magnetic stand, then washed with 300 *μ*L of 1× TWB (Tween Washing Buffer; 5 mM Tris-HCl pH 7.5; 0.5 mM EDTA; 1 M NaCl; 0.05% Tween 20). The beads were resuspended in 300 *μ*L of 2× Binding Buffer (10 mM Tris-HCl pH 7.5, 1 mM EDTA; 2 M NaCl), the sonicated DNA was added (diluted to a final volume of 300 *μ*L if necessary), and the beads were incubated for ≥15 minutes at room temperature on a rotator. After separation on a magnetic stand, the beads were washed with 300 *μ*L of 1× TWB, and heated at 55 °C in a Thermomixer with shaking for 2 minutes. After removal of the supernatant on a magnetic stand, the TWB wash and 55 °C incubation were repeated.

Final libraries were prepared on beads using the NEB-Next Ultra II DNA Library Prep Kit (NEB, #E7645) as follows. End repair was carried out by resuspending beads in 50 *μ*L 1× EB buffer, and adding 3 *μ*L NEB Ultra End Repair Enzyme and 7 *μ*L NEB Ultra End Repair Enzyme, followed by incubation at 20 °C for 30 minutes (in a Thermomixer, with shaking at 1,000 rpm) and then at 65 °C for 30 minutes.

Adapters were ligated to DNA fragments by adding 30 *μ*L Blunt Ligation mix, 1 *μ*L Ligation Enhancer and 2.5 *μ*L NEB Adapter, incubating at 20 °C for 20 minutes, adding 3 *μ*L USER enzyme, and incubating at 37°C for 15 minutes (in a Thermomixer, with shaking at 1,000 rpm).

Beads were then separated on a magnetic stand, and washed with 300 *μ*L TWB for 2 minutes at 55 °C, 1000 rpm in a Thermomixer. After separation on a magnetic stand, beads were washed in 100 *μ*L 0.1 × TE buffer, then resuspended in 15 *μ*L 0.1 × TE buffer, and heated at 98 °C for 10 minutes.

For PCR, 5 *μ*L of each of the i5 and i7 NEB Next sequencing adapters were added together with 25 *μ*L 2× NEB Ultra PCR Mater Mix. PCR was carried out with a 98 °C incubation for 30 seconds and 12 cycles of 98 °C for 10 seconds, 65 °C for 30 seconds, and 72 °C for 1 minute, followed by incubation at 72 °C for 5 minutes.

Beads were separated on a magnetic stand, and the supernatant was cleaned up using 1.8× AMPure XP beads.

Libraries were sequenced in a paired-end format on a Illumina NextSeq instrument using NextSeq 500/550 high output kits (2×36 cycles).

### Hi-C experiments

Hi-C was carried out using the previously described *in situ*procedure^46^as follows:

*B. natans* cells were first crosslinked using 37% formaldehyde (Sigma) at a final concentration of 1% for 15 minutes at room temperature. Formaldehyde was then quenched using 2.5 M Glycine at a final concentration of 0.25 M. Cells were subsequently centrifuged at 2,000 *g* for 5 minutes, washed once in 1×PBS, and stored at −80 °C.

Cell lysis was initiated by incubation with 250 *μ*L of cold Hi-C Lysis Buffer (10 mM Tris-HCl pH 8.0, 10 mM NaCl, 0.2% Igepal CA630) on ice for 15 minutes, followed by centrifugation at 2,500 *g* for 5 minutes, a wash with 500 *μ*L of cold Hi-C Lysis Buffer, and centrifugation at 2,500 *g* for 5 minutes. The pellet was the resuspended in 50 *μ*L of 0.5% SDS and incubated at 62 °C for 10 minutes. SDS was quenched by adding 145 *μ*L of H_2_O and 25 *μ*L of 10% Triton X-100 and incubating at 37 °C for 15 minutes.

Restriction digestion was carried out by adding 25 *μ*L of 10 ×NEBuffer 2 and 100 U of the MluCI restriction enzyme (NEB, R0538) and incubating for ≥2 hours at 37°C in a Thermomixer at 900 rpm. The MluCI restriction enzyme was chosen as more suitable for the highly AT-rich nucleomorph genome. The reaction was then incubated at 62 °C for 20 minutes in order to inactivate the restriction enzyme.

Fragment ends were filled in by adding 37.5 *μ*L of 0.4 mM biotin-14-dATP (ThermoFisher Scientific, # 19524-016), 1.5 *μ*L each of 10 mM dCTP, dGTP and dTTP, and 8 *μ*L of 5U/*μ*L DNA Polymerase I Large (Klenow) Fragment (NEB M0210). The reaction was the incubated at 37°C in a Thermomixer at 900 rpm for 45 minutes.

Fragment end ligation was carried out by adding 663 *μ*L H_2_O, 120 *μ*L 10×NEB T4 DNA ligase buffer (NEB B0202), 100 *μ*L of 10% Triton X-100, 12 *μ*L of 10 mg/mL Bovine Serum Albumin (100× BSA, NEB), 5 *μ*L of 400 U/*μ*L T4 DNA Ligase (NEB M0202), and incubating at room temperature for ≥4 hours with rotation.

Nuclei were then pelleted by centrifugation at 2,000 *g*for 5 minutes; the pellet was resuspended in 200 *μ*L ChIP Elution Buffer (1% SDS, 0.1 M NaHCO_3_), Proteinase K was added, and incubated at 65 °C overnight to reverse crosslinks.

After addition of 600 *μ*L 1× TE buffer, DNA was sheared using a Covaris E220 instrument. DNA was then purified using the MinElute PCR Purificaiton Kit (Qiagen #28006), with elution in a total volume of 300 *μ*L 1 × EB buffer.

For streptavidin pulldown of biotin-labeled DNA, 150 *μ*L of 10 mg/mL Dynabeads MyOne Streptavidin T1 beads (Life Technologies, 65602) were separated on a magnetic stand, then washed with 400 *μ*L of 1× TWB (Tween Washing Buffer; 5 mM Tris-HCl pH 7.5; 0.5 mM EDTA; 1 M NaCl; 0.05% Tween 20). The beads were resuspended in 300 *μ*L of 2×Binding Buffer (10 mM Tris-HCl pH 7.5, 1 mM EDTA; 2 M NaCl), the sonicated DNA was added, and the beads were incubated for ≥15 minutes at room temperature on a rotator. After separation on a magnetic stand, the beads were washed with 600 *μ*L of 1× TWB, and heated at 55 °C in a Thermomixer with shaking for 2 minutes. After removal of the supernatant on a magnetic stand, the TWB wash and 55 °C incubation were repeated.

Final libraries were prepared on beads using the NEB-Next Ultra II DNA Library Prep Kit (NEB, #E7645) as follows. End repair was carried out by resuspending beads in 50 *μ*L 1× EB buffer, and adding 3 *μ*L NEB Ultra End Repair Enzyme and 7 *μ*L NEB Ultra End Repair Enzyme, followed by incubation at 20 °C for 30 minutes and then at 65 °C for 30 minutes.

Adapters were ligated to DNA fragments by adding 30 *μ*L Blunt Ligation mix, 1 *μ*L Ligation Enhancer and 2.5 *μ*L NEB Adapter, incubating at 20 °C for 20 minutes, adding 3 *μ*L USER enzyme, and incubating at 37°C for 15 minutes.

Beads were then separated on a magnetic stand, and washed with 600 *μ*L TWB for 2 minutes at 55 °C, 1000 rpm in a Thermomixer. After separation on a magnetic stand, beads were washed in 100 *μ*L 0.1 × TE buffer, then resuspended in 16 *μ*L 0.1 × TE buffer, and heated at 98 °C for 10 minutes.

For PCR, 5 *μ*L of each of the i5 and i7 NEB Next sequencing adapters were added together with 25 *μ*L 2× NEB Ultra PCR Mater Mix. PCR was carried out with a 98 °C incubation for 30 seconds and 12 cycles of 98 °C for 10 seconds, 65 °C for 30 seconds, and 72 °C for 1 minute, followed by incubation at 72 °C for 5 minutes.

Beads were separated on a magnetic stand, and the supernatant was cleaned up using 1.8× AMPure XP beads.

Libraries were sequenced in a paired-end format on a Illumina NextSeq instrument using NextSeq 500/550 high output kits (either 2×75 or 2×36 cycles).

### ATAC-seq data processing

Demultiplexed FASTQ files were mapped to the v1.0 assembly for *Bigelowiella natans* CCMP2755 (with the nucleomorph sequence added) as 2×36mers using Bowtie^47^ with the following settings: `-v 2 -k 2 -m 1 --best --strata -X 1000`. Duplicate reads were removed using `picard-tools` (version 1.99). Reads mapping to the plastid, mitochondrion and the nucleomoprh were filtered out for the analysis of accessibility in the nuclear genome.

Browser tracks generation, fragment length estimation, TSS enrichment calculations, and other analyses were carried out using custom-written Python scripts (https://github.com/georgimarinov/GeorgiScripts).

For the purpose of the analysis of rDNA arrays in nucleomorphs, alignments were carried out with unlimited multimappers with the following settings: `-v 2 -a --best --strata -X 1000`. Normalization of multimappers was performed as previously described^48^.

### ATAC-seq peak calling

Peak calling was carried out using version 2.1.0 of MACS2^29^ with default settings.

### Analysis of positioned nucleosomes

Positioned nucleosomes along the whole nucleomorph genome and in the ±500 bp regions around annotated TSSs in the nucleus were identified using NucleoATAC^31^ as follows. We used the low resolution nucleosome calling program `nucleoatac occ` with default parameters that requires ATAC-seq data and genomic windows of interest, and returns a list of nucleosome positions based on the distribution of ATAC-seq fragment lengths centered at these positions. To cover the whole nucleomorph genome, sliding windows of 1 kbp in steps of 500 bp were taken as inputs, and redundant nucleosome positions were eventually discarded. For nuclear TSSs, 1-kbp windows centered at the TSSs were used as inputs. V plots were made by aggregating unique-mapping ATAC-seq reads centered around the positioned nucleosomes, and mapping the density of fragment sizes versus fragment center locations relative to the positioned nucleosomes as previously described^31,32^.

### KAS-seq data processing

Demultiplexed FASTQ files were mapped to the v1.0 assembly for *Bigelowiella natans* CCMP2755 (with the nucleomorph sequence added) as 2×36mers using Bowtie^47^ with the following settings: `-v 2 -k 2 -m 1 --best --strata -X 1000`. Duplicate reads were removed using `picard-tools` (version 1.99).

Browser tracks generation, fragment length estimation, TSS enrichment calculations, and other analyses were carried out using custom-written Python scripts (https://github.com/georgimarinov/GeorgiScripts).

For the analysis of rDNA arrays in nucleomorphs, alignments were carried out with unlimited multimappers with the following settings: `-v 2 -a --best --strata -X 1000`. Normalization of multimappers was performed as previously described^48^.

### Hi-C data processing and assembly scaffolding

As an initial step, Hi-C sequencing reads were processed against the previously published *B. natans* assembly^41^ using the Juicer pipeline^49^ for analyzing Hi-C datasets (version 1.8.9 of Juicer Tools).

The resulting Hi-C matrices were then used as input to the 3D DNA pipeline^40^ for automated scaffolding with the following parameters:
`--editor-coarse-resolution 5000 --editor-coarse-region 5000 --polisher-input-size 100000 --polisher-coarse-resolution 1000`
`--polisher-coarse-region 300000`
`--splitter-input-size 100000`
`--splitter-coarse-resolution 5000`
`--splitter-coarse-region 300000 --sort-output`
`--build-gapped-map -r 10 -i 5000`.

Manual correction of obvious assembly and scaffolding errors was then carried out using Juicebox^49^.

After finalizing the scaffolding, Hi-C reads were reprocessed against the new assembly using the Juicer pipeline.

## Author contributions

G.K.M. conceptualized the study, performed cell culture, ATAC-seq, KAS-seq and Hi-C experiments and analyzed data. X.C. carried out nucleosome positioning analysis. T.W. and C.H. provided key reagents. W.J.G., A.K., and A.R.G. supervised the study. G.K.M. wrote the manuscript with input from all authors.

## Acknowledgements

This work was supported by NIH grants (P50HG007735, RO1 HG008140, U19AI057266 and UM1HG009442 to W.J.G., 1UM1HG009436 to W.J.G. and A.K., 1DP2OD022870-01 and 1U01HG009431 to A.K.), the Rita Allen Foundation (to W.J.G.), the Baxter Foundation Faculty Scholar Grant, and the Human Frontiers Science Program grant RGY006S (to W.J.G). W.J.G. is a Chan Zuckerberg Biohub investigator and acknowledges grants 2017-174468 and 2018-182817 from the Chan Zuckerberg Initiative. Fellowship support provided by the Stanford School of Medicine Dean’s Fellowship (G.K.M.). This work is also supported by NSF-IOS EDGE Award 1645164 to A.R.G.

The authors would like to thank Alexandro E. Trevino and members of the Greenleaf, Kundaje, Grossman and Pringle laboratories for helpful discussion and suggestions regarding this work.

## Data Availability

Data associated with this manuscript have been submitted to GEO under accession number GSE181751.

## Code Availability

Custom code used to process the data is available at https://github.com/georgimarinov/GeorgiScripts and https://github.com/chenxy19/nucleomorph.

## Supplementary Materials

### Supplementary Figures

**Supplementary Figure 1:**
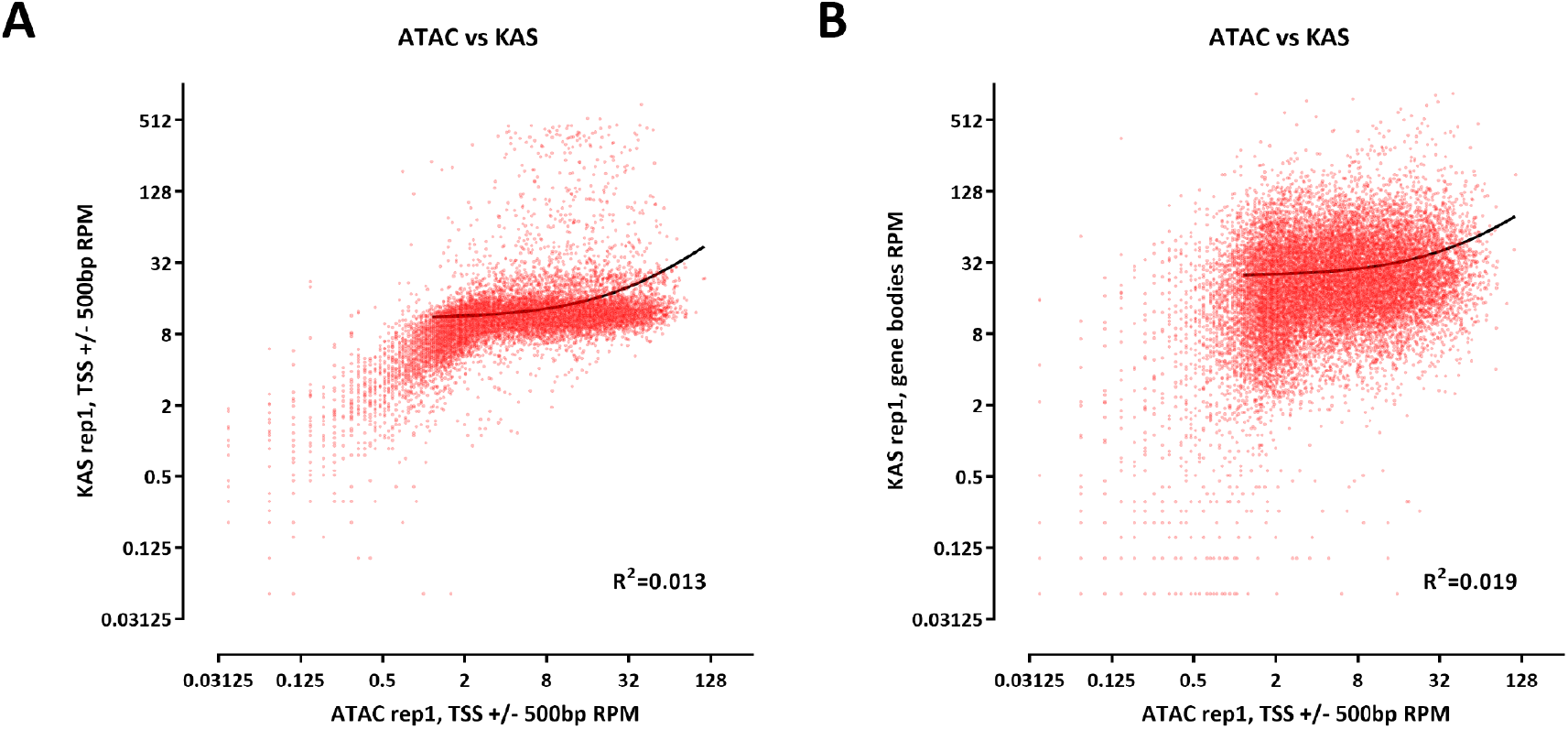
Relationship between chromatin accessibility and active transcription as measured by KAS-seq in the *B. natans* nuclear genome. (A) Correlation between ATAC-seq signal over promoters and KAS-seq signal over promoters. (B) Correlation between ATAC-seq signal over promoters and KAS-seq signal over gene bodies.

**Supplementary Figure 2:**
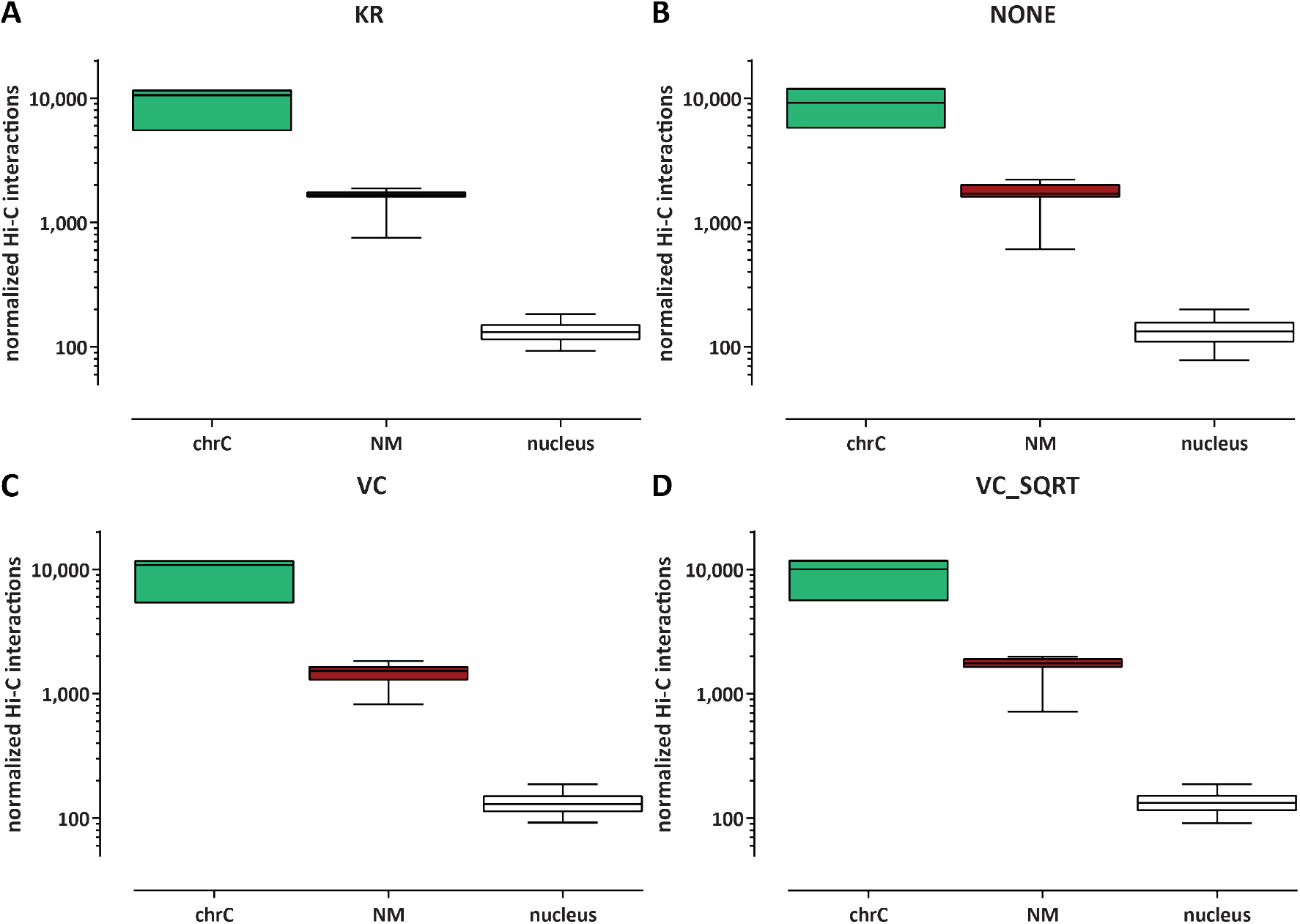
Impact of different Hi-C normalization methods on the quantification of Hi-C *trans* contacts between different compartments. (A) KR normalization (B) No normalization (C) Coverage normalization (VC) (D) Coverage normalization (VC_SQRT)

